# Nutrient availability shifts the biosynthetic potential of soil-derived microbial communities

**DOI:** 10.1101/2021.09.02.458721

**Authors:** Marc G Chevrette, Bradley Himes, Camila Carlos-Shanley

## Abstract

Secondary metabolites produced by microorganisms are the main source of antimicrobials other pharmaceutical drugs. Soil microbes have been the primary discovery source for these secondary metabolites, often producing complex organic compounds with specific biological activities. Research suggests that secondary metabolism broadly shapes microbial ecological interactions, but little is known about the factors that shape the abundance, distribution, and diversity of biosynthetic gene clusters in the context of microbial communities. In this study, we investigate the role of nutrient availability on the abundance of biosynthetic gene clusters in soil-derived microbial consortia. We found that soil microbial consortia enriched in medium with 150 mg/L of glucose and 200 mg/L of trehalose (here defined as high sugar) had more biosynthetic gene cluster and higher inhibitory activity than soil microbial consortia enriched in medium with 15 mg/L of glucose + 20 mg/L of trehalose (here defined as low sugar). Our results demonstrate that laboratory microbial communities are a promising tool to study ecology of specialized metabolites.

## Introduction

The chemical products of microbial secondary metabolism (also called natural products) modulate interactions within and between species and are thus a major means through which the microbial world communicates [1]. Secondary metabolites have had an enormous impact on modern medicine: they are the main source of antimicrobials used to treat infections, they have been used as therapeutics for cancer and other important human diseases, and as immunosuppressants that enable life-saving transplantation surgeries [2]. Soil microbes have been the primary discovery source for these secondary metabolites, often producing complex organic compounds with specific biological activities [3]. The enzymes that assemble microbial natural products are encoded by genes located in biosynthetic gene clusters (BGCs). While it is widely hypothesized that secondary metabolism broadly shapes microbial ecological interactions, little is known about the factors that shape the abundance, distribution, and diversity of biosynthetic gene clusters in the context of microbial communities [4].

Some studies have attempted to show the gross differences in biosynthetic potential between the microbial communities of different soil biomes, but these studies have limited extrapolative value, and little is known of how environmental factors can contribute to enrichment for secondary metabolism on finer scales. Comparisons between United States soil communities from New Hampshire and Arizona suggest that the arid desert soils of Arizona may harbor more antagonistic, inhibitory compounds than the forest soils of New Hampshire [5]. They observed a diversity of Nonribosomal peptide synthetases (NRPSs) and polyketide synthases (PKSs) domains in arid soils when compared with forest soils. One hypothesis is that this is due to the harsh, nutrient poor conditions of the soil that may lead to increased pressures on nutrient acquisition and/or other means of competition. Even within soils sourced from the same rhizospheres, biosynthetic capacity sees shifts depending on soil depth [3]. BGCs and their producing organisms are found in almost every known microbial niche and covary with some environmental [6]. However, these studies are often limited to correlative descriptions subject to sampling biases and systematic assessments with sufficient experimental control remain lacking.

Many bacteria dedicate very large portions of their genomes to BGCs, sometimes in excess of 25% of all genetic material [7], that is often maintained vertically over evolutionary timescales [8, 9], implying that they are important in their natural settings [8], yet under controlled laboratory conditions the producers of secondary metabolites often do not express their BGCs, therefore not producing their chemical products. A growing body of evidence is showing that a battery of different culture conditions and/or perturbations is needed to the production of different elicit secondary metabolite [10]. Recently, Hurley et al. [11], observed that the taxonomy and inhibitory profile of the bacteria isolated from four United States soil samples used within the Tiny Earth project is strongly affect by the selective media used. Potato dextrose agar (PDA) enriched for strains that inhibited *Acinetobacter baylyi* and *Pseudomonas putida*, while tryptic soy agar (TSA) enriched for *Erwinia carotavora* inhibiting strains. Understanding the link between secondary metabolisms and nutrient availability has fundamental implications across microbial ecology, including the ecology of antagonism, community maintenance, invasion, niche construction, and niche defense. The aim of this study was to evaluate if carbon source availability can affect the biosynthetic potential of enriched microbial communities.

## Material and Methods

### Sample Collection and community enrichment

Soil sample was collected from the Purgatory Creek Natural Area in San Marcos, Texas (GPS coordinates 29.882029, −97.982068). The sample came from small field off of a trail surrounded by a wooded area. Once the sample was taken back to the lab it was passed through a sieve washed with 70% ethanol to filter out larger particles, the remaining sample was weighed and then suspended in sterile PBS at a concentration of 1.5 mL of PBS per 0.1 g of soil sample. Once the suspension had been made it was then used to inoculate six Erlenmeyer flasks containing 50 mL of M63 Minimal Media with a 5 mL/L concentration of SPV-4 Trace elements and a 1 mL/L concentration of MgSO4. The three low sugar group flasks (Media LS) contained 15 mg/L of glucose and 20 mg/L of trehalose while the high sugar group flasks (Media HS) contained 150 mg/L of glucose and 200 mg/L of trehalose. Media LS concentrations of glucose and trehalose were based on Jenkins et al. (2017) [12]. The inoculated flasks were cultured at 30°C and 225 rpm, the OD600 of the culture was measured every 24 hours to monitor growth stage, the samples grown in the lower nutrient flasks were called PCA1-3 while those in the higher nutrient flaks were called PCB1-3. Once stationary phase had been reached samples of each of the flasks were gathered and then centrifuged, the cell pellets were frozen at −80°C while the supernatant was kept refrigerated to be used with certain microbes in a 96-well growth assay to determine antimicrobial potential.

### Inhibition assays

To determine if the metabolites produced in the six cultures at stationary phase had antimicrobial properties an inhibitory assay was developed that measured the inhibitory activity of the metabolites at different concentrations against a panel of selected bacteria. The bacterial target panel selected was compromised of bacterial species *Pseudomonas fluorescens* (ATCC13525), *Klebsiella pneumoniae* (ATCC23357), *Bacillus Subtilis* (ATCC6051), and *Salmonella typhimirium* LT2 (Nickerson-Arizona State). Target bacteria were grown on a 96 well plate containing 10μL sterile Thermo Scientific Iso-Sensitest broth + 90μL microbial community supernatant in quadruplicate for each treatment. Growth was measured using optical density at 625 nm via a BioTek plate reader and Gen5 software. Readings were taken at timepoints 0, 24 and 48 hours after inoculation, plates were grown at 37°C and 26°C depending on which bacterial species was being tested (*P. fluorescens* and *B. subtilis* at 26°C, *K. pnuemoniae* and *S. typhimirium* at 37°C). Controls were frown with 90μL of sterile M63 minimal medium with carbon source + 10μL sterile Thermo Scientific Iso-Sensitest. The optical density data measured from the control was then used in comparison to the data from each respective community supernatant to determine if the supernatants contained metabolites with inhibitory effects since the growth recorded from the control plates was in the presence of no antimicrobial metabolites

### Metagenomic DNA Extraction, Sequencing and Analysis

DNA extraction and purification were performed for all six samples using the QIAamp BiOstic Bacteremia DNA Kit and protocol (QIAGEN). The purified sample DNA was sequenced at the Microbial Genome Sequencing Center (MiGS) core facility in Pittsburgh on the Illumina NextSeq platform. Sequence reads were filtered and trimmed using the default settings of fastp (Chen et al., 2018). Filtered reads were taxonomically classified using the Kaiju software using the NCBI BLAST nr+euk database [13]. Co-occurrence network was built using the SparCC [14] program implements in the MicrobiomeAnalyst platform [15] based on the Spearman correlation between genus distribution across the datasets. Metagenomes were assembled using SPAdes 3.13.0 [16]. Biosynthetic gene clusters (BGCs) were annotated with antiSMASH 5.0 using the default settings [17]. Contigs with BGCs were taxonomic classified using a Last Common Ancestor (LCA) approach implemented in the Contig Annotation Tool (CAT) [18]. The short reads of the metagenome datasets used in this study were deposited in the NCBI Short Read Archive (SRA) accession numbers from SRR15633242-SRR15633247.

### Results and Discussion

In this study, soil microbial communities were enriched in minimal medium containing either 15 mg/L of glucose + 20 mg/L of trehalose (here defined as low sugar, LS) or 150 mg/L of glucose and 200 mg/L of trehalose (here defined as high sugar, HS). The supernatant of the microbial communities enriched on the high sugar (HS) medium was able to inhibit the growth of *B. subtilis* (t=5.413, df=3, p=0.012), *K. pneumoniae* (t=3.846, df=3, p=0.031), *P. fluorescens* (t=9.565, df=3, p=0.002) and *S. typhimurium* (t=5.249, df=3, p=0.13). While the supernatant of the microbial communities enriched on the low sugar (LS) medium was able to inhibit the growth of *K. pneumoniae* (t=4.65, df=3, p=0.019) and *S. typhimurium* (t=4.801, df=3, p=0.017) (Figure 1). Some studies have found that high concentrations of nutrient, such as glucose, inhibit the production of secondary metabolites in some bacteria [19]. However, it seems that different carbon sources can have different effects on the production of secondary metabolites in different bacteria [20]. To our knowledge, our study is the first to evaluate the effect of glucose and trehalose concentrations on the inhibitory activity of an undefined microbial consortia. While the approach used in this study does not enable the identification of the major producers of inhibitory molecules it allows to study how nutrient availability shapes interspecies interactions.

**Figure 1.**
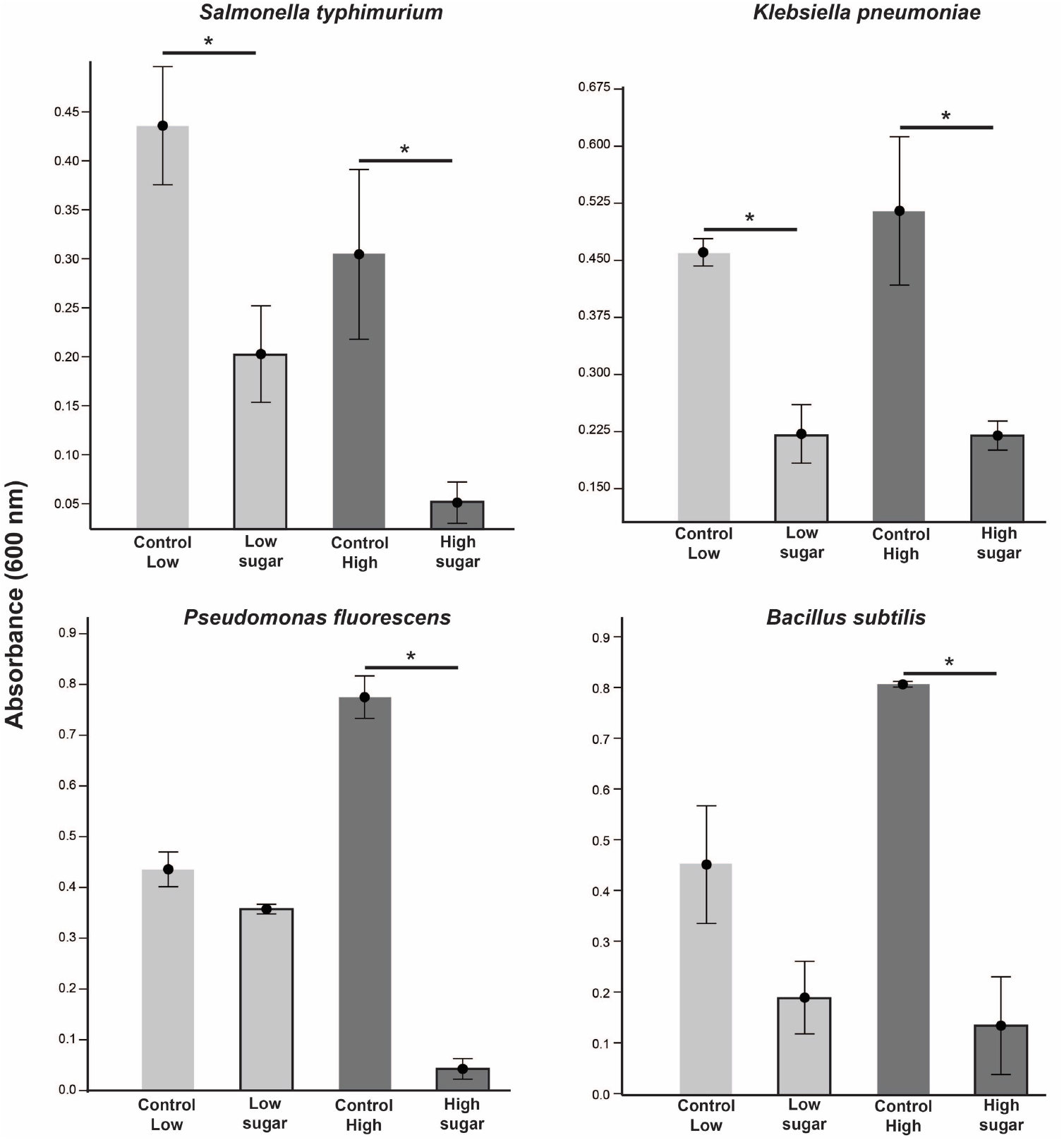
Effect of spent media of soil-derived microbial communities on the growth of *Salmonella typhimurium*, *Klebsiella pneumoniae*, *Pseudomonas fluorescens* and *Bacillus subtilis*. Absorbance at 600 nm was measured after 48h and results shown as average of three experiments. Asterisks (*) refers to significant comparisons (T-test, p<0.05) between spent media of soil-derived microbial communities growing on low and high sugar medium and the fresh medium control.

The number of reads in the metagenomic datasets after quality filtering and trimming ranged from 9 to 10 million. Metagenomic assemblies ranged from 51 to 390 Mb (Table 1). Taxonomic classification of reads showed that the HS microbial communities were dominated by fungi, while LS microbial communities were dominated by bacteria (Figure 2a). Previous studies have evaluated the effect of nutrient availability on microbial community diversity, abundance, and composition. Experiment with soils amended with glucose found that concentrations higher than 8mg C/g of soil favored the growth of fungi over bacteria, which the authors attribute to difference in optimal osmotic potential [21].

**Table 1.** Metagenome information.

| Treatment | Sample | # of million reads | # of Contigs | Assembly length in Mb | GC % | # of BGCs | Total length of BGC's in Mb | % of base pairs in BGCs |
| --- | --- | --- | --- | --- | --- | --- | --- | --- |
| Low sugar | PCA1 | 9,7 | 1,092,758 | 390.8 | 0.4975 | 64 | 0.431052 | 0.11027591 |
|  | PCA2 | 9,8 | 759,937 | 326.9 | 0.4535 | 65 | 0.703316 | 0.2150899 |
|  | PCA3 | 9,2 | 81,499 | 51.4 | 0.5892 | 51 | 0.355935 | 0.69244013 |
| High sugar | PCB1 | 10,7 | 3,98,017 | 237.9 | 0.4856 | 139 | 3.543708 | 1.48934092 |
|  | PCB2 | 10,7 | 621,134 | 325.1 | 0.4785 | 122 | 1.336774 | 0.41122777 |
|  | PCB3 | 10,7 | 373,335 | 236.8 | 0.4902 | 168 | 1.54994 | 0.65444703 |

BGCs= Biosynthetic gene clusters

**Figure 2.**
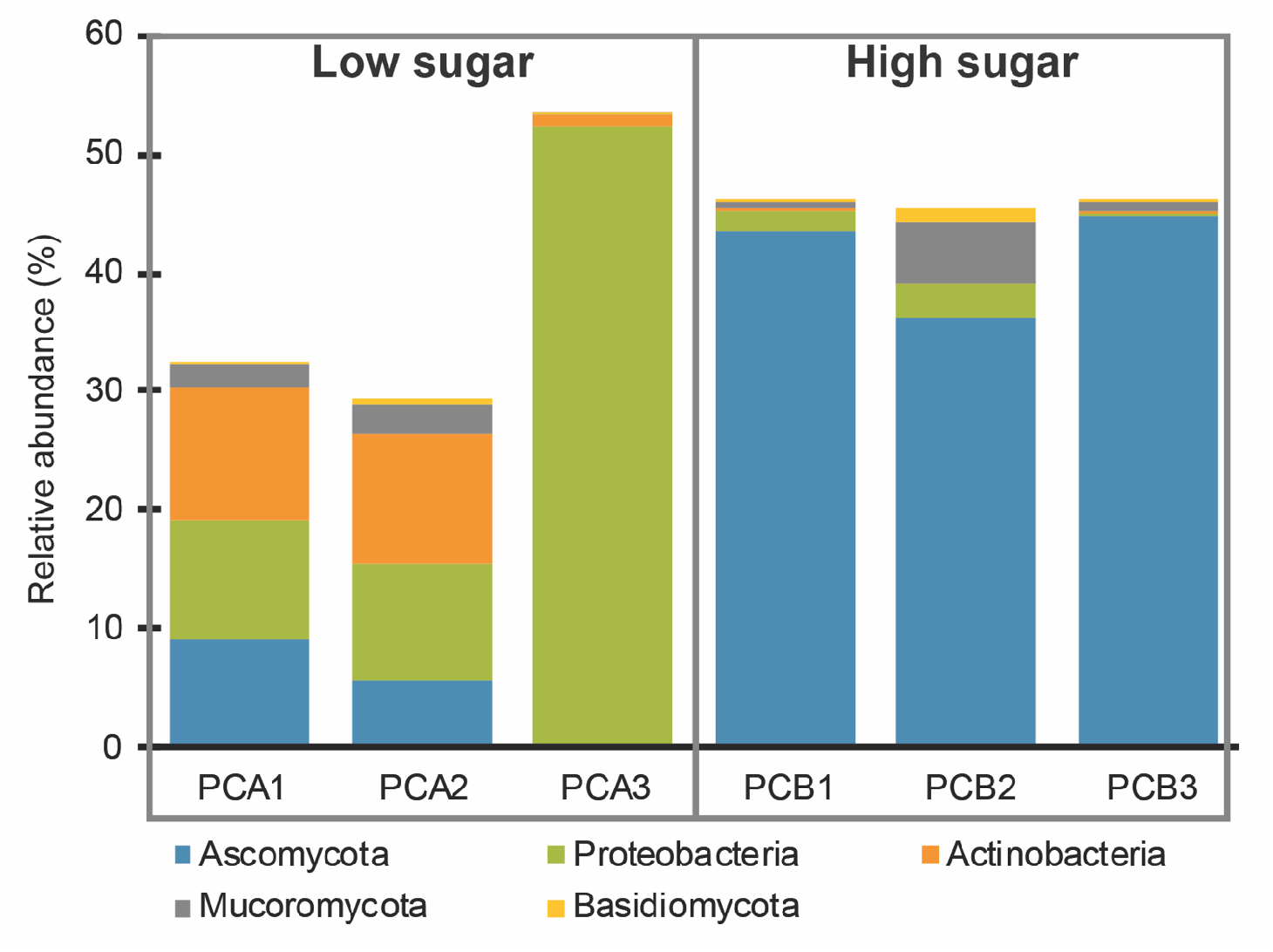
Taxonomic profile of metagenomic reads. Relative abundance of the top 5 phyla. Samples PCA1-3 are biological replicates of the low sugar treatment and PCB1-3 are biological replicates of the high sugar treatment.

Functional annotation of the contigs in Clusters of Orthologous Genes (COG) categories revealed that genes in the “Chromatin structure and dynamics” (t=8.214, df=4, p=0.007) and “Secondary metabolites biosynthesis, transport, and catabolism” (t=5.915, df=4, p=0.018) categories were overrepresented in the HS microbial communities, while genes in “Signal transduction mechanisms” (t=4.156, df=4, p=0.021) were overrepresented in the LS microbial communities (Figure 3). Most biosynthetic gene clusters (BGCs) were overrepresented in the HS microbial communities. Type I Polyketide Synthetase (T1PKS) and Non-Ribosomal Peptide Synthetase (NPRS) biosynthetic gene clusters were more abundant in the HS communities, while BGCs encoding bacteriocins and siderophores were more abundant in the LS communities (Table 2). Correlation analysis of microbial genera in the communities revealed many negative correlations between the two most abundant fungal genera *Trichoderma* and *Fusarium* (both belonging to the Sordariomycetes class) and bacteria belonging to the Enterobacteriaceae family, such as *Salmonella* (Figure 4a). Most of BGCs in the HS microbial communities were classified to the Sordariomycetes taxonomic class, while most of BGCs in the LS microbial communities were classified to the Actinobacteria taxonomic class (Figure 4b).

**Figure 3.**
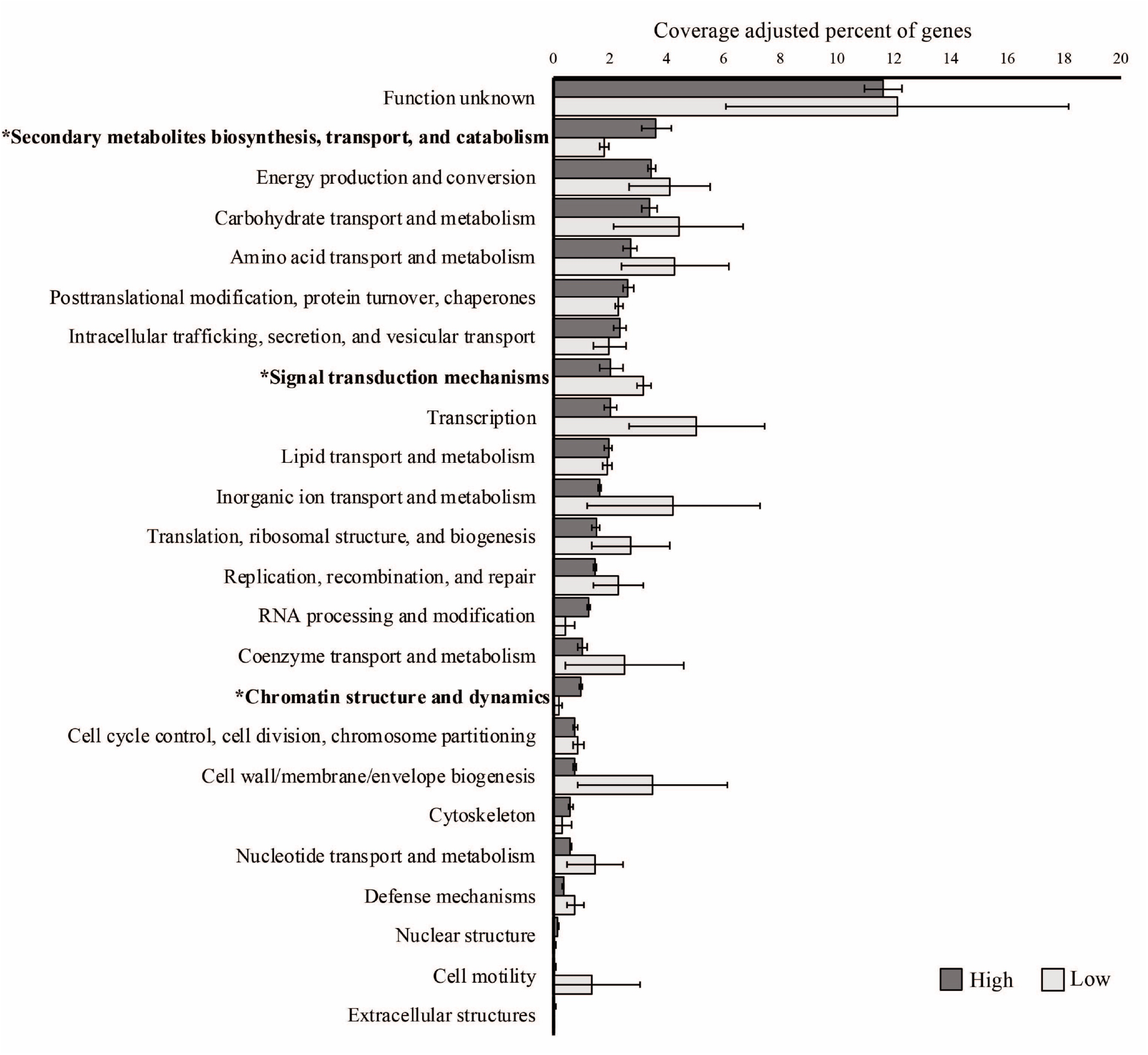
Percentage of genes in COG categories in the low and high sugar microbial communities (coverage adjusted). Values of the categories marked with an asterisk (*) were significantly different between the treatments (T-test, p-values<0.05).

**Table 2.** Distribution of biosynthetic gene cluster (BGC) classes across the metagenomic datasets.

| BGC class | Low sugar |  |  | High sugar |  |  |
| --- | --- | --- | --- | --- | --- | --- |
|  | PCA1 | PCA2 | PCA3 | PCB1 | PCB2 | PCB3 |
| NRPS | 15 | 13 | 12 | 58 | 55 | 91 |
| NRPS-like | 5 | 5 | 6 | 23 | 32 | 22 |
| terpene | 21 | 16 | 8 | 16 | 13 | 17 |
| T1PKS | 1 | 1 | 1 | 32 | 15 | 27 |
| bacteriocin | 7 | 9 | 6 | 0 | 0 | 0 |
| siderophore | 3 | 6 | 3 | 0 | 2 | 0 |
| NRPS\|T1PKS | 0 | 1 | 0 | 4 | 3 | 3 |
| other | 12 | 14 | 15 | 6 | 2 | 8 |

**Figure 4.**
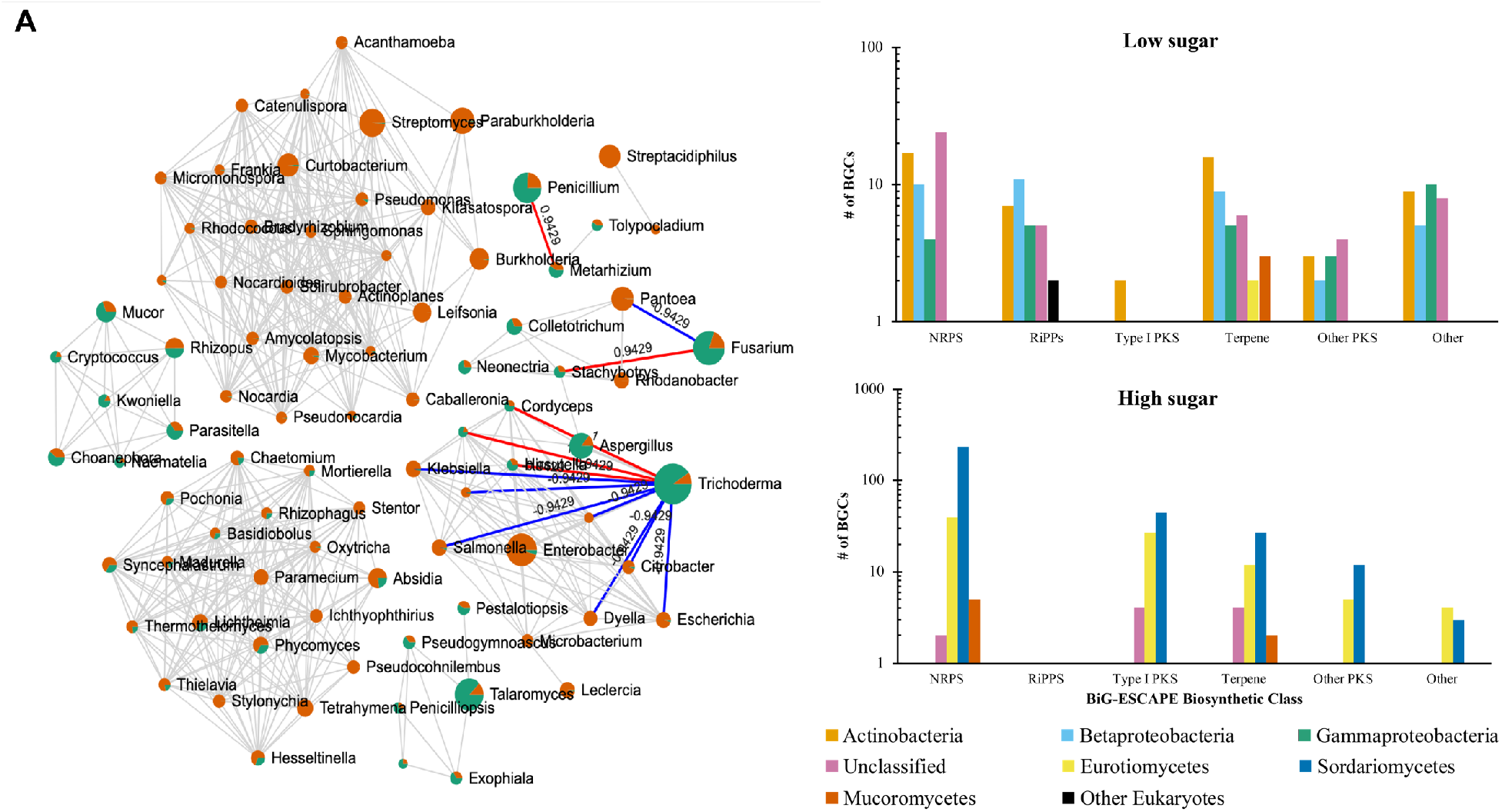
A. Co-occurrence network where nodes are microbial genera and edges are Spearman correlation values > 0.8 and with p-value < 0.01. Node size is proportional to genus abundance and the average distribution of each genus in the low sugar treatment is shown in orange and in the high sugar treatment is shown in green. Negative (blue) and positive (red) correlations of *Trichoderma*, *Fusarium* and *Penicillium* are highlighted. B. Distribution of biosynthetic gene clusters (BGCs) across microbial class. Contigs with BGCs were taxonomic classified using a Last Common Ancestor (LCA) approach. NRPS = Nonribosomal peptide synthetase; PKS = polyketide synthase; RiPP = Ribosomally synthesized and post-translationally modified peptide;

Microbial communities with more BGCs were showed more inhibitory activity towards the bacterial pathogens used in this study. Traditionally, inhibition assays and other screening assays for antimicrobial activity are performed with axenic cultures, and the use of mixed cultures for antibiotic discovery is still in its early days [22, 23]. It has been hypothesized that co-culture can activate silent biosynthetic gene clusters and facilitate the discovery of new natural products [24, 25]. In this study we demonstrate that enriched microbial communities derived from environments with complex microbial communities, such as soil and feces, could be screened for the production of novel antibacterial compounds.

## Conclusion

In conclusion, in this study we show that laboratory microbial communities are a promising tool to study ecology of specialized metabolites. Future studies involving fraction libraries, metatranscriptomic, and metabolomic approaches will contribute to our understanding of the environmental and nutritional conditions that are favorable for production and/or selection of novel secondary metabolite.

## Acknowledgments

Support for M.G.C. provided by grant 2020-67012-31772 (accession 1022881) from the USDA National Institute of Food and Agriculture.

## Author contributions

C.C.-S. and B.H. designed the experiments. B.H. performed the experiments. M.G.C. and C.C.-S. performed bioinformatic and statistical analyses. M.G.C., B.H. and C.C.-S. wrote and reviewed the manuscript.

